# The structural complexity of the Gammaproteobacteria flagellar motor is related to the type of its torque-generating stators

**DOI:** 10.1101/369397

**Authors:** Mohammed Kaplan, Debnath Ghosal, Poorna Subramanian, Catherine M. Oikonomou, Andreas Kjær, Sahand Pirbadian, Davi R. Ortega, Mohamed Y. El-Naggar, Grant J. Jensen

## Abstract

The bacterial flagellar motor is a cell-envelope-embedded macromolecular machine that functions as a propeller to move the cell. Rather than being an invariant machine, the flagellar motor exhibits significant variability between species, allowing bacteria to adapt to, and thrive in, a wide range of environments. For instance, different torque-generating stator modules allow motors to operate in conditions with different pH and sodium concentrations and some motors are adapted to drive motility in high-viscosity environments. How such diversity evolved is unknown. Here we use electron cryo-tomography to determine the *in situ* macromolecular structures of the flagellar motors of three Gammaproteobacteria species: *Legionella pneumophila*, *Pseudomonas aeruginosa*, and *Shewanella oneidensis* MR-1, providing the first views of intact motors with dual stator systems. Complementing our imaging with bioinformatics analysis, we find a correlation between the stator system of the motor and its structural complexity. Motors with a single H^+^-driven stator system have only the core P- and L-rings in their periplasm; those with dual H^+^-driven stator systems have an extra component elaborating their P-ring; and motors with Na^+^- (or dual Na^+^-H^+^)- driven stator systems have additional rings surrounding both their P- and L-rings. Our results suggest an evolution of structural complexity that may have enabled pathogenic bacteria like *L. pneumophila* and *P. aeruginosa* to colonize higher-viscosity environments in animal hosts.

## Introduction

The bacterial flagellum is a macromolecular machine that transforms the movement of ions (H^+^ or Na^+^) across the cell membrane into a mechanical torque to move the bacterial cell through its environment[1]. In general, the flagellum consists of a cell-envelope-embedded motor, a hook which acts as a universal joint and a long propeller-like filament[2,3]. The motor can rotate the filament in either a counterclockwise or clockwise direction. For cells with a single flagellum this drives the cell forward or backward; for peritrichous cells this results in "run" or "tumble" movements. Flagella can also exhibit more complex behavior; it was recently reported that the *Shewanella putrefaciens* flagellum can wrap around the cell to mediate a screw-like motion that allows the cell to escape narrow traps[4]. Besides their role in motility, bacterial flagella participate in other vital activities of the cell such as biofilm formation[5]. Moreover, the virulence of many human pathogens depends directly on their flagella, with flagellated strains of *Pseudomonas aeruginosa* and *Legionella pneumophila* causing more serious infections with higher mortality rates[6,7]. *P. aeruginosa* lacking fully-assembled flagella cause no mortality and are 75% less likely to cause pneumonia in mice[6].

The best-studied flagellar motor, in *Salmonella enterica*, consists of several sub-complexes, which we will describe in order from the inside out. On the cytoplasmic side are the inner-membrane-embedded MS ring (formed by the protein FliF) and the C-ring (aka the switch complex, formed by FliN, FliM and FliG). The C-ring encircles a type III secretion system (T3SS) export apparatus (FliH, FliI, FliJ, FlhA, FlhB, FliP, FliQ and FliR). Spanning the space from the inner membrane to the peptidoglycan cell wall is the ion channel (called the stator), a complex of two proteins (MotA and MotB) with 4:2 stoichiometry[8,9]. The interaction between the stator and the C-ring (FliG) generates the torque required to drive the flagellum. The MS ring is coupled to the extracellular hook (FlgE) through the rod (FlgB, FlgC, FlgF and FlgG). The rod is further surrounded by two other rings: the P- (peptidoglycan, FlgI) and the L- (lipopolysaccharide, FlgH) rings which act as bushings during rod rotation. Extending from the hook is the filament (FliC) which is many micrometers in length. In addition to these components, the assembly of the whole flagellar motor is a highly synchronized process that requires a plethora of additional chaperones and capping proteins[10–12].

Recently, the development of electron cryo-tomography (ECT)[13,14] has allowed the determination of the complete structures of flagellar motors in their cellular milieu at macromolecular (~5 nm) resolution. ECT studies of many different bacterial species have revealed that while the core structure described above is conserved, the flagellar motor has evolved many species-specific adaptations to different environmental conditions[15–21]. For example, extra periplasmic rings were found to elaborate the canonical P- and L-rings in the motor of the Gammaproteobacteria *Vibrio* species. These rings are called the T-ring (MotX and Y) and H-ring (FlgO, P and T)[20]. Unlike the *S. enterica* motor described above, which is driven by H^+^ ions, the motors of *Vibrio* and other marine bacteria employ different stators (PomA and PomB) which utilize Na^+^. These Na^+^-dependent stators generate higher torque (~2,200 pN) than H^+^-dependent stators (~1,200 pN), driving the motor at higher speeds (up to 1,700 Hz compared to ~300 Hz in H^+^-driven motors)[22].

Most flagellated bacteria use a single stator system – either H^+^-driven or Na^+^-driven, depending on their environment. Some species, however, such as *Vibrio alginolyticus*, use two distinct types of motors to move in different environments: a polar Na^+^-driven flagellum and lateral H^+^-driven flagella. Still other species employ dual stator systems with a single flagellar motor, conferring an advantage for bacteria that experience a range of environments (see [23] and references therein). For example, *P. aeruginosa* employs a dual H^+^-driven stator system (MotAB and MotCD). While the MotAB system is sufficient to move the cell in a liquid environment[24], MotCD is necessary to allow the cell to move in more viscous conditions[25]. *Shewanella oneidensis* MR-1 combines both Na^+^- and H^+^-dependent stators in a single motor, enabling the bacterium to move efficiently under conditions of different pH and Na^+^ concentration[26]. How these more elaborate motors may have evolved remains an open question.

Here, we used ECT to determine the first *in situ* structures of three Gammaproteobacteria flagellar motors with dual stator systems: in *L. pneumophila*, *P. aeruginosa* and *S. oneidensis* MR-1. *L. pneumophila* and *P. aeruginosa* have dual H^+^-dependent stator systems and *S. oneidensis* has a dual Na^*+*^-H^+^-dependent stator. This imaging, along with bioinformatics analysis, shows a correlation between the structural complexity of the motor and its stator system, suggesting a possible evolutionary pathway.

## Results and Discussion

To determine the structures of the flagellar motors of *L. pneumophila*, *P. aeruginosa,* and *S. oneidensis* we imaged intact cells of each species in a hydrated frozen state using ECT. We identified clearly visible flagellar motors in the tomographic reconstructions and performed sub-tomogram averaging to enhance the signal-to-noise ratio, generating a 3D average of the motor of each species at macromolecular resolution (Fig. 1 and S1). While all three motors shared the conserved core structure of the flagellar motor, they exhibited different periplasmic decorations surrounding this conserved core. While the *S. oneidensis* and *P. aeruginosa* averages showed clear densities corresponding to the stators (Fig. 1 E, F, K and L, orange density), none were visible in the *L. pneumophila* average, suggesting that they were more variable, or dynamic. Interestingly, we observed a novel feature in the *S. oneidensis* motor: an extra ring outside the outer membrane (Fig. 1 A-F, purple density). This structure is reminiscent of the O-ring (outer membrane ring) described recently in the sheathed flagellum of *Vibrio alginolyticus*[17]. However, while the *V. alginolyticus* O-ring was associated with a 90° bend in the outer membrane, no such outer membrane bend was seen in the unsheathed *S. oneidensis* flagellum, so the function of this structure remains mysterious.

**Figure 1:**
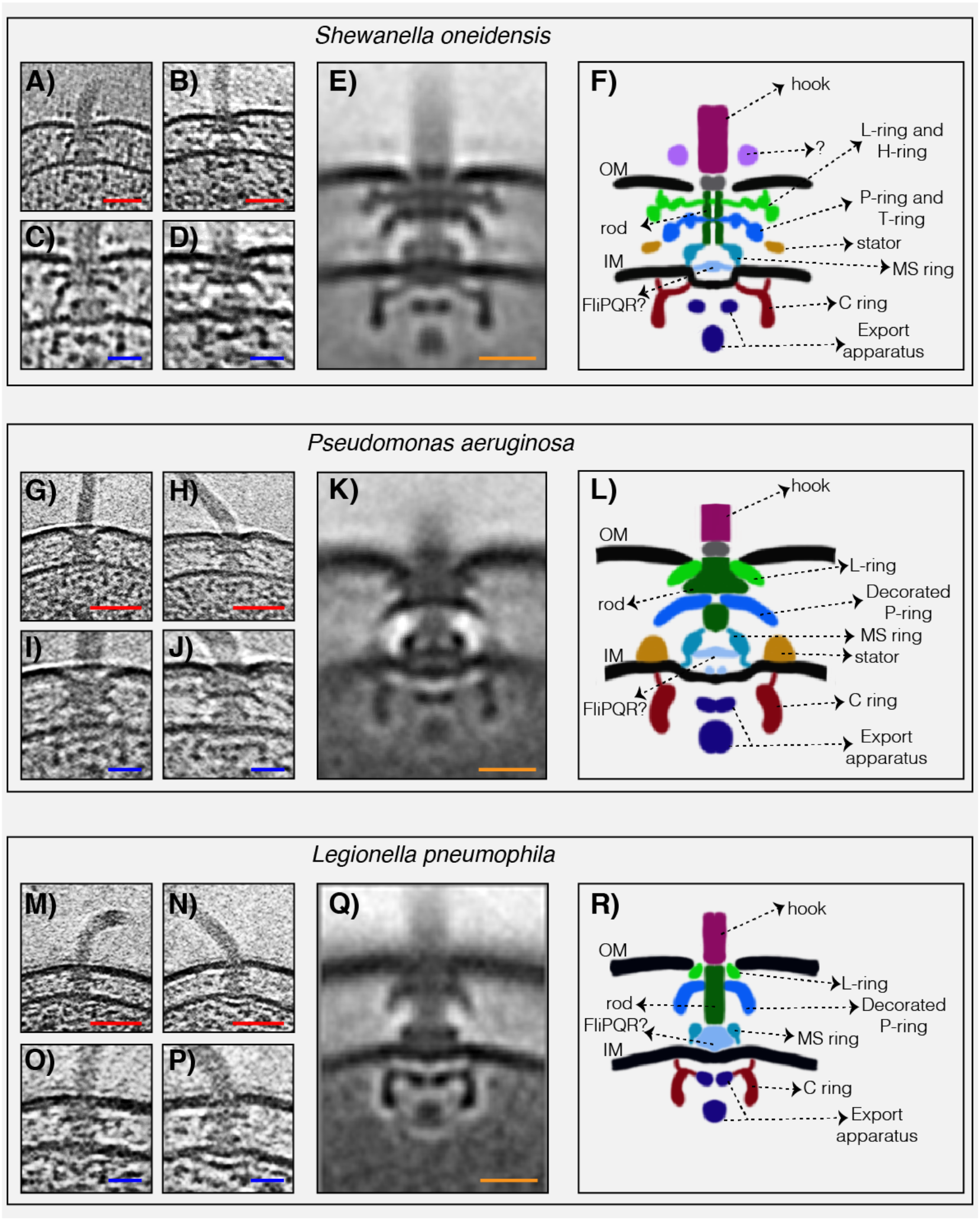
The structures of three dual-stator Gammaproteobacteria flagellar motors revealed by ECT. **A & B)** slices through *Shewanella oneidensis* MR-1 electron cryo-tomograms showing single polar flagella. **C & D)** zoomed-in views of the slices shown in **A** and **B** highlighting the flagellar motors. **E)** central slice through a sub-tomogram average of the *S. oneidensis* MR-1 flagellar motor. **F)** schematic representation of the sub-tomogram average shown in **E** with the major parts of the motor labeled. **G-L)** flagellar motor of *Pseudomonas aeruginosa*. Panels follow the same scheme as in **A-F** above. **M-R)** flagellar motor of *Legionella pneumophila*. Panels follow the same scheme as above. Scale bars 50 nm (red) and 20 nm (blue and orange).

The most striking difference between the three motor structures was the L- and P-rings, which were highly elaborated in *S. oneidensis*. The *P. aeruginosa* and *L. pneumophila* motors lacked additional rings associated with the L-ring, but showed smaller elaborations of their P-rings. To determine whether flagellar motor structure correlates with motor type, we compared our three new ECT structures with those of the five previously-published Gammaproteobacteria motors (Fig. 2). Two motors (*Escherichia coli* and *S. enterica*) have a single H^+^-driven stator system, two motors have dual H^+^-dependent stator systems (*P. aeruginosa* and *L. pneumophila*), three motors have Na^+^-driven systems (the three *Vibrio* species) and one motor has a dual Na^+^-H^+^-driven system (*S. oneidensis*). Interestingly, we found that motors with similar stator type also shared similar structural characteristics. While the two motors with a single H^+^-dependent stator system did not show any periplasmic elaborations beyond the conserved flagellar core, the dual H^+^-dependent stator systems had an extra ring surrounding their P-ring, with no embellishment of the L-ring. The Na^+^-dependent motors of the *Vibrio spp.*, together with the Na^+^-H^+^-dependent motor of *S. oneidensis* have extra components surrounding both their P- and L- rings. In *Vibrio*, these extra periplasmic rings are known as the T-ring (surrounding the P-ring and formed by the MotX and MotY proteins) and the H-ring (surrounding the L-ring and consisting of the FlgO, FlgP and FlgT proteins). The presence of the T- and H-rings was suggested to be specific to the Na^+^-driven *Vibrio* motors[20] with the FlgT protein required for the formation of both rings[27].

**Figure 2:**
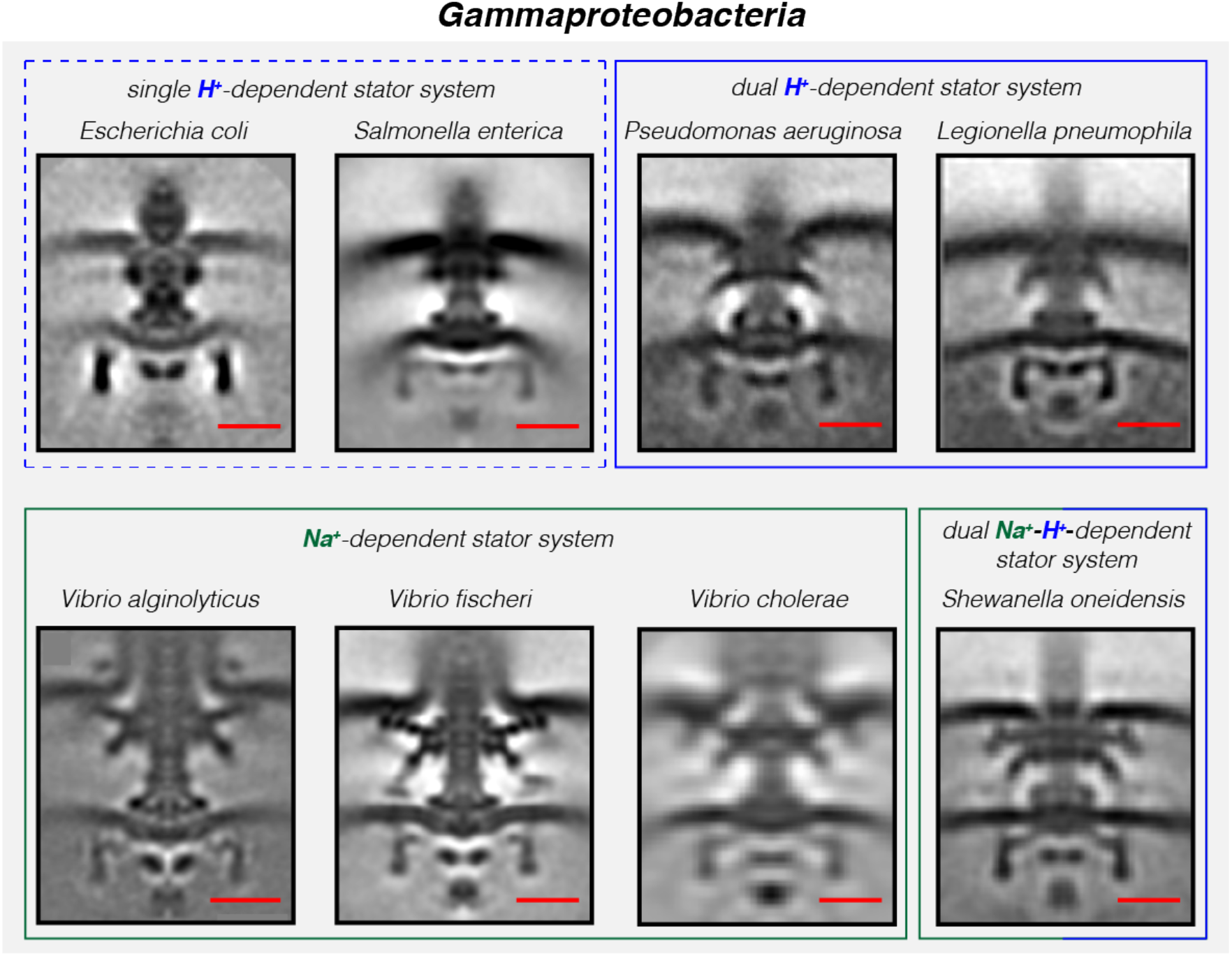
Compilation of all Gammaproteobacteria flagellar motors imaged to date by ECT. Central slices of sub-tomogram averages are shown for the eight Gammaproteobacteria flagellar motors revealed by ECT, including the three structures solved in this study (*P. aeruginosa, L. pneumophila* and *S. oneidensis*). The motors are classified based on their stator system: single H^+^-driven (dashed blue box), dual H^+^-driven (blue box), Na^+^-driven (green box) or dual Na^+^-H^*+*^-driven (green-blue box). Scale bars are 20 nm.

Previous studies showed that MotX and MotY are important for flagellar rotation in *S. oneidensis* but it was not known whether they form part of the motor or not[28]. Similarly, bioinformatics analysis and biochemical studies showed that MotY is involved in the function of the *P. aeruginosa* motor, but the structural basis of this role was not known[24]. We therefore performed a bioinformatics search for candidate homologs of MotX, MotY, FlgO, FlgP and FlgT in the genomes of *P. aeruginosa*, *L. pneumophila* and *S. oneidensis* to examine whether there is a correlation between the presence of homologous genes and the extra periplasmic rings observed in the ECT structures. While we found candidates for all five proteins constituting the T- and H-rings in *S. oneidensis* as previously suggested[29], only MotY candidates were found in *L. pneumophila* and *P. aeruginosa* (Table S1). This is in accordance with our ECT structures, which showed that *L. pneumophila* and *P. aeruginosa* motors have a ring surrounding only their P-rings while the *S. oneidensis* motor has rings surrounding both the P- and L-rings. These rings are likely T- and H-rings, respectively, as in *Vibrio*. The lack of candidate MotX homologs in the genomes of *L. pneumophila* and *P. aeruginosa* (Table S1) is consistent with their lack of PomB, the component of the Na^+^- dependent stator with which MotX interacts. Interestingly, the absence of candidates for FlgT in the *L. pneumophila* and *P. aeruginosa* genomes suggests that it may not be required for the recruitment of MotY as in *Vibrio* species.

To see whether these correlations hold more broadly, we expanded our bioinformatics analysis to additional species of Gammaproteobacteria. We examined the genomes of species with single H^+^-driven stator systems (Table S2), dual H^+^-driven stator systems (Table S3) and Na^+^-driven stator systems (Table S4). Interestingly, we identified a second species, *Colwellia psychrerythraea* 34H, with a single motor and candidates for both PomAB (Na^+^-driven) and MotAB (H^+^-driven) stator systems, similar to *S. oneidensis* MR-1 (Table S5). In all species we examined, we observed the same pattern: (i) genomes of species with single H^+^-driven stator systems lacked homologs of H- or T- ring components; (ii) genomes of species with Na^+^ stator systems contained homologs of all H- and T-ring components, and (iii) genomes of species with dual H^+^-driven stator systems contained candidate homologs only for the T-ring component MotY. The sole exception to this rule was *Chromohalobacter salexigens* DSM 3043, which contained a homolog of FlgO in addition to MotY. None of the eight species with dual H^+^-driven stator systems we examined contained a homolog of FlgT, further suggesting that it is not essential for MotY stabilization in this group.

Together, our results from ECT imaging of flagellar motors *in situ* and bioinformatics analysis reveal a correlation between the structural complexity of the flagellar motor of Gammaproteobacteria and the type of its torque-generating unit, the stator (summarized in Fig. 3). Low-speed motors with single H^+^-stator systems have only the P- and L-ring, while high-speed motors using Na^+^ have two extra periplasmic rings, the T- and H-rings. Unexpectedly, we find that motors with dual H^+^-driven stator systems represent a hybrid structure between the two, elaborating their P-rings with one of the five components of the T- and H-rings, MotY. This extra MotY ring might help to stabilize the motor under conditions of increased load, as in the viscous environment of the pulmonary system encountered by *L. pneumophila* and *P. aeruginosa*. These results therefore suggest an evolutionary pathway in which these pathogenic Gammaproteobacteria species could have borrowed a motor stabilization strategy from related Na^+^-driven motors to allow them to colonize animal hosts.

**Figure 3:**
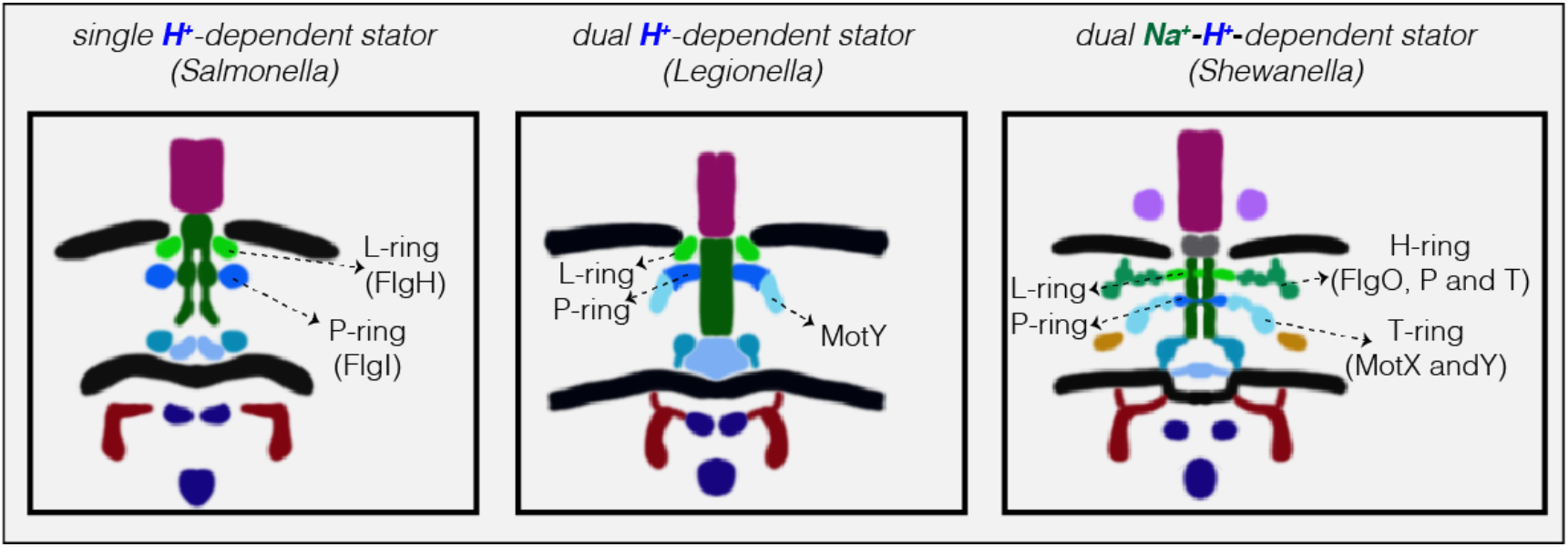
Models showing correlation between structural complexity of the flagellar motor and its stator type. Flagellar motors with single H^+^-driven stator systems (e.g. *Salmonella*) have P- and L-rings alone. Motors with dual H^+^-driven stator systems have an extra ring surrounding the P-ring formed by the MotY protein alone. Motors with Na^+^-driven motors have two periplasmic rings, the T-ring (MotX and MotY) and H-ring (FlgO, FlgP and FlgT), decorating the P- and L-rings respectively. Note that the boundaries between the P- and L-rings and their decorations are tentative in these schematics.

## Acknowledgements

This work is supported by the National Institutes of Health (NIH, grant R01 AI127401 to G.J.J.). M.K. is supported by a postdoctoral Rubicon fellowship from De Nederlandse Organisatie voor Wetenschappelijk Onderzoek (NWO). S.P. and M.Y.E.-N. are supported by the Air Force Office of Scientific Research Presidential Early Career Award for Scientists and Engineers (FA955014-1-0294, to M.Y.E.-N.).

## Supporting information

**Figure S1:**
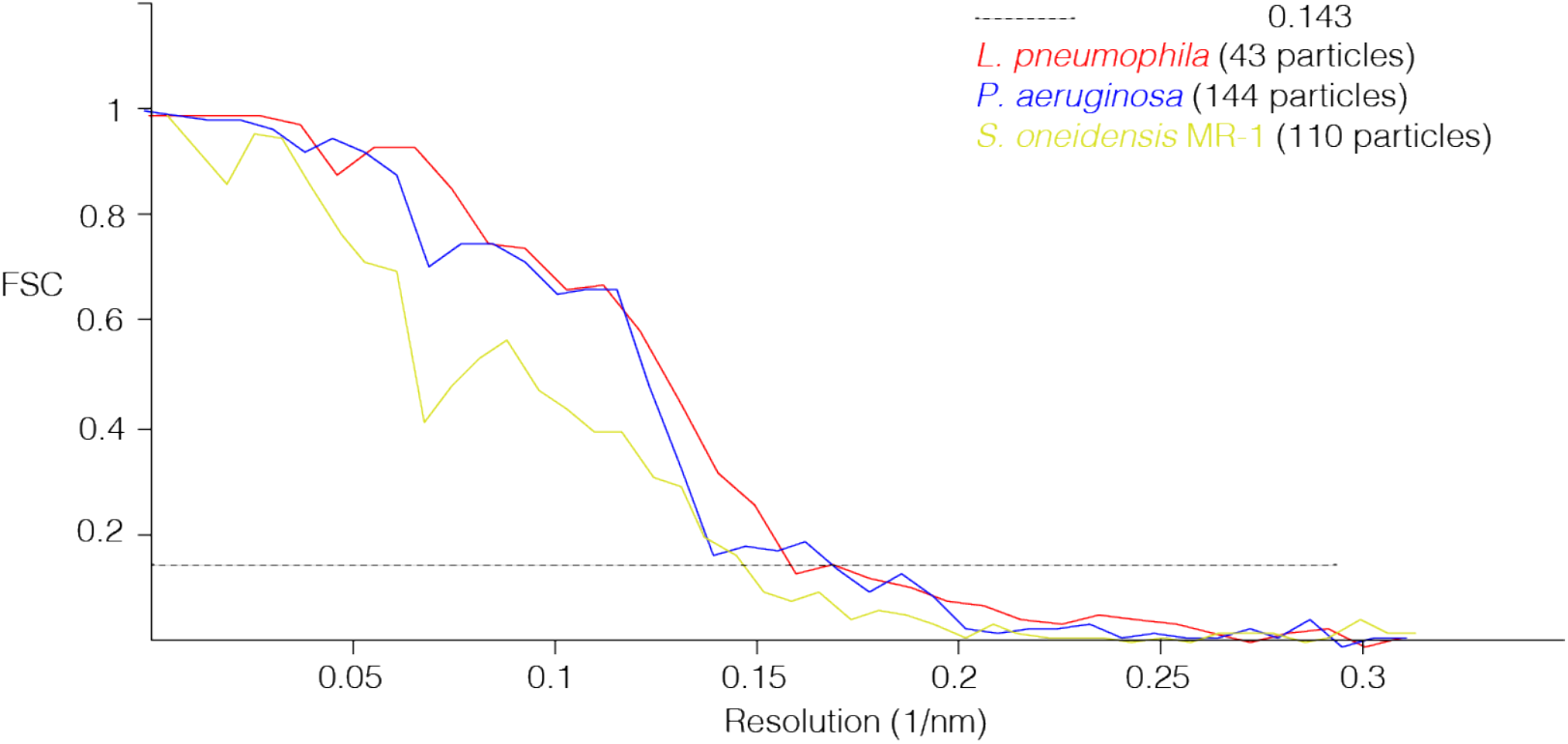
Gold-standard FSC curves of sub-tomogram averages. Resolutions at a 0.143 cutoff (dashed line) are: *L. pneumophila*, 6.4 nm; *P. aeruginosa*, 5.9 nm; *S. oneidensis* MR-1, 6.9 nm.

**Table S1.**
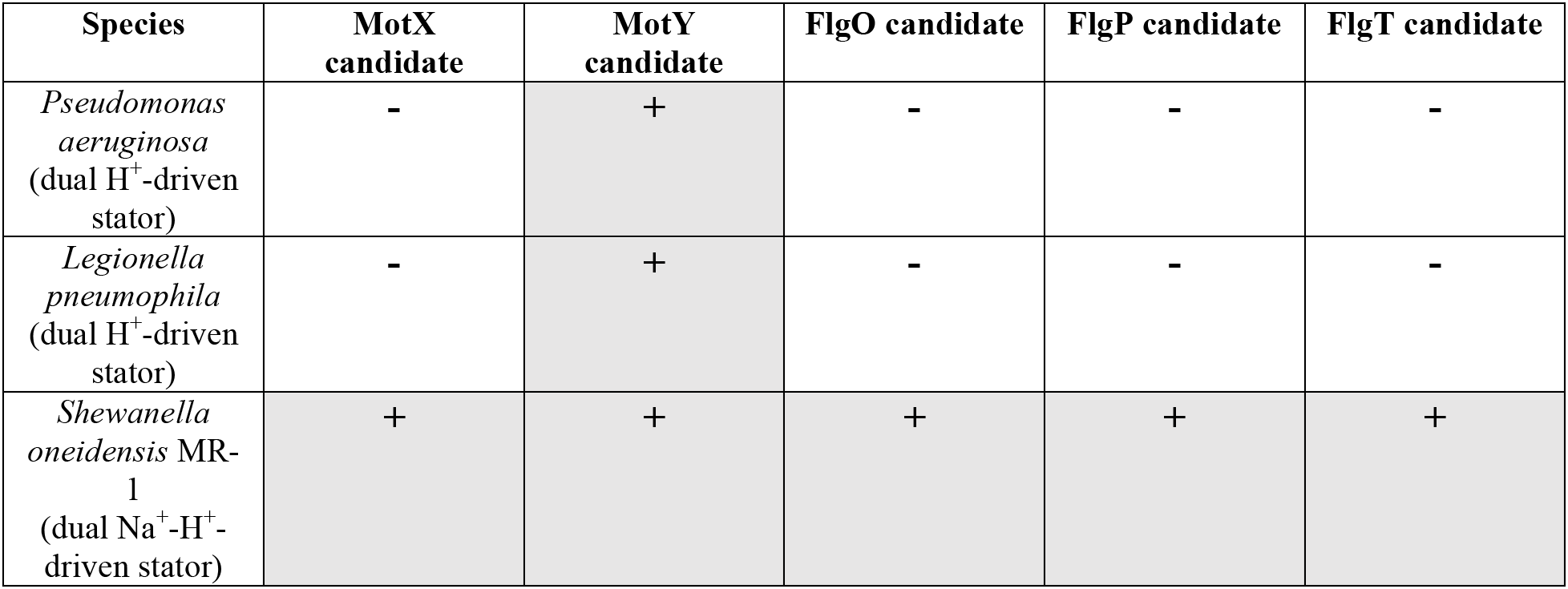
Candidate homologs of H- and T-ring components in species imaged in this study.

| Species | MotX candidate | MotY candidate | FlgO candidate | FlgP candidate | FlgT candidate |
| --- | --- | --- | --- | --- | --- |
| *Pseudomonas aeruginosa*<br>(dual H<sup>+</sup>-driven stator) | - | + | - | - | - |
| *Legionella pneumophila*<br>(dual H<sup>+</sup>-driven stator) | - | + | - | - | - |
| *Shewanella oneidensis* MR-1<br>(dual Na<sup>+</sup>-H<sup>+</sup>-driven stator) | + | + | + | + | + |

**Table S2.** Candidate homologs of H- and T-ring components in single H^+^-dependent stator systems.

| Species | MotX candidate | MotY candidate | FlgO candidate | FlgP candidate | FlgT candidate |
| --- | --- | --- | --- | --- | --- |
| *Escherichia coli* | - | - | - | - | - |
| *Salmonella enterica* | - | - | - | - | - |
| *Sodalis glossinidius* | - | - | - | - | - |

**Table S3.** Candidate homologs of H- and T-ring components in dual H^+^-dependent stator systems.

| Species | MotX candidate | MotY candidate | FlgO candidate | FlgP candidate | FlgT candidate |
| --- | --- | --- | --- | --- | --- |
| *Azotobacter vinelandii* DJ | - | + | - | - | - |
| *Cellvibrio japonicus* Ueda107 | - | + | - | - | - |
| *Chromohalobacter salexigens* DSM 3043 | - | + | + | - | - |
| *Pseudomonas entomophila* | - | + | - | - | - |
| *Saccharophagus degradans* 2-40 | - | + | - | - | - |
| *Xanthomonas campestris* pv. *campestris* | - | + | - | - | - |
| *Teredinibacter turnerae* T7901 | - | + | - | - | - |
| *Pseudomonas putida* | - | + | - | - | - |

**Table S4.** Candidate homologs of H- and T-ring components in Na^+^-dependent stator systems.

| Species | MotX candidate | MotY candidate | FlgO candidate | FlgP candidate | FlgT candidate |
| --- | --- | --- | --- | --- | --- |
| *Colwellia psychrerythraea* 34H | + | + | + | + | + |
| *Vibrio alginolyticus* | + | + | + | + | + |
| *Vibrio fischeri* | + | + | + | + | + |

**Table S5.** Candidate homologs of stator system components in *Colwellia psychrerythraea* 34H. *E*-values of BLAST results are shown for each candidate locus (name in parentheses).

| Species | MotA candidate | MotB candidate | PomA candidate | PomB candidate |
| --- | --- | --- | --- | --- |
| *Colwellia psychrerythraea* 34H | 8e-10<br>(CPS_1524) | 4e-11<br>(CPS_1525) | 2e-124<br>(CPS_1092) | 4e-129<br>(CPS_1093) |

**Table S6.** Raw Blast results for all species in Tables S1-S5. E-values are shown. For *E*-values exceeding the cutoff, the top hit is listed in parentheses.

| Species | MotX candidate | MotY candidate | FlgO candidate | FlgP candidate | FlgT candidate |
| --- | --- | --- | --- | --- | --- |
| *Azotobacter vinelandii* DJ | 1e-06 | 8e-14<br>(Avin_48650) | 0.032 | 0.32 | 4.2 |
| *Cellvibrio japonicas* Ueda107 | 0.59 | 9e-28<br>(CJA_2588) | 0.72 | 0.29 | 6e-09 |
| *Chromohalobacter salexigens* DSM 3043 | 0.014 | 6e-13<br>(Csal_3309) | 9e-16<br>(Csal_2511) | 2.8 | 2.7 |
| *Pseudomonas entomophila* | 3e-05 | 2e-31<br>(PSEEN1209) | 1.6 | 0.32 | 0.099 |
| *Saccharophagus degradans* 2-40 | 6e-06 | 1e-37<br>(Sde_2427) | 0.24 | 2.1 | 1.2 |
| *Xanthomonas campestris* pv. *campestris* | 0.019 | 1e-13<br>(XCC1436) | 3 | 4.5 | 0.021 |
| *Teredinibacter turnerae* T7901 | 0.7 | 3e-35<br>(TERTU_3000) | 1.4 | 0.061 | 1 |
| *Pseudomonas putida* | 0.005 | 7e-31<br>(PP_1087) | 1 | 0.77 | 0.063 |
| *Legionella pneumophila* | 1e-08 | 3e-35<br>(lpg2962) | 0.87 | 0.11 | 2.2 |
| *Pseudomonas aeruginosa* | 3e-05 | 2e-37<br>(PA3526) | 0.047 | 0.63 | 0.19 |
| *Escherichia coli* | 9e-06 | 2e-09 | 0.19 | 0.84 | 0.19 |
| *Salmonella enterica* | 5e-07 | 5e-09 | 0.34 | 4.9 | 0.82 |
| *Sodalis glossinidius* | 4.4 | 1e-07 | 0.88 | 0.27 | 2.2 |
| *Colwellia psychrerythraea*<br>34H | 2e-63<br>(CPS_4618) | 1e-73<br>(CPS_3471) | 2e-59<br>(CPS_1469) | 6e-28<br>(CPS_1470) | 5e-38<br>(CPS_1468) |
| *Shewanella oneidensis* MR-1 | 2e-46<br>(SO_3936) | 2e-80<br>(SO_2754) | 2e-19<br>(SO_3257) | 6e-31<br>(SO_3256) | 3e-36<br>(SO_3258) |

## Materials and Methods

### Strains and growth conditions

*Legionella pneumophila* (strain Lp02) cells were grown on plates of ACES [N-(2-acetamido)-2-aminoethanesulfonic acid]-buffered charcoal yeast extract agar (CYE) or in ACES-buffered yeast extract broth (AYE) with 100 μg/ml thymidine. Ferric nitrate and cysteine hydrochloride were added to the media. For ECT experiments, cells were harvested in early stationary phase.

*Shewanella oneidensis* MR-1 cells belonging to the strains listed in Table S7 were used in this study. They were grown using one of the following methods: Luria–Bertani (LB) broth culture, chemostat, the batch culture method or in a perfusion flow imaging platform. Detailed descriptions of these methods can be found in[30]. Briefly, in the chemostat method, 5 mL of a stationary-phase overnight LB culture was injected into a continuous flow bioreactor containing an operating liquid volume of 1 L of a defined medium[31], while dissolved oxygen tension (DOT) was maintained at 20%. After 20 h, and as the culture reached stationary phase, continuous flow of the defined medium[31] was started with a dilution rate of 0.05 h^−1^ while DOT was still maintained at 20%. After 48 h of aerobic growth under continuous flow conditions, the DOT was manually reduced to 0%. O_2_ served as the sole terminal electron acceptor throughout the experiment. pH was maintained at 7.0, temperature at 30 °C, and agitation at 200 rpm. Either 24 or 40 hours after DOT reached 0%, samples were taken from the chemostat for ECT imaging.

In the batch culture method, 200 μL of an overnight LB culture of *S. oneidensis* cells was added to each of two sealed and autoclaved serum bottles containing 60 mL of a defined medium[31]. One of the two bottles acted as a control and was not used for imaging. To this control bottle, 5 μM resazurin was added to indicate the O_2_ levels in the medium. The bottles were then placed in an incubator at 30 °C, with shaking at 150 rpm until the color due to resazurin in the control bottle completely faded, indicating anaerobic conditions. At this point, samples were taken for ECT imaging from the bottle that did not contain resazurin.

For the perfusion flow imaging experiments, *S. oneidensis* cells were grown overnight in LB broth at 30 °C to an OD_600_ of 2.4–2.8 and washed twice in a defined medium[31]. A glow-discharged, carbon-coated, R2/2, Au NH2 London finder Quantifoil EM was glued to a 43 mm × 50 mm no. 1 glass coverslip using waterproof silicone glue (General Electric Company) and let dry for ~30 min. Using a vacuum line, the perfusion chamber (model VC-LFR-25; C&L Instruments) was sealed against the grid-attached glass coverslip. A total of ^~^10 mL of the washed culture was injected into the chamber slowly to allow cells to settle on the grid surface, followed by a flow of sterile defined medium from an inverted serum bottle through a bubble trap (model 006BT-HF; Omnifit) into the perfusion chamber inlet. Subsequently, the flow of medium was stopped and the perfusion chamber was opened under sterile medium. The grid was then detached from the coverslip by scraping off the silicone glue at the grid edges using a 22-gauge needle and rinsed by transferring three times in deionized water, before imaging by ECT.

Samples were also prepared from an aerobic *S. oneidensis* LB culture grown at 30 °C to an OD_600_ of 2.4–2.8.

*Pseudomonas aeruginosa* PAO1 cells were first grown on LB plates at 37 °C overnight. Subsequently, cells were inoculated into 5 ml MOPS [(3-(*N*-morpholino) propanesulfonic acid)] Minimal Media Limited Nitrogen and grown for ~ 24 hours at 30 °C.

**Table S7.**
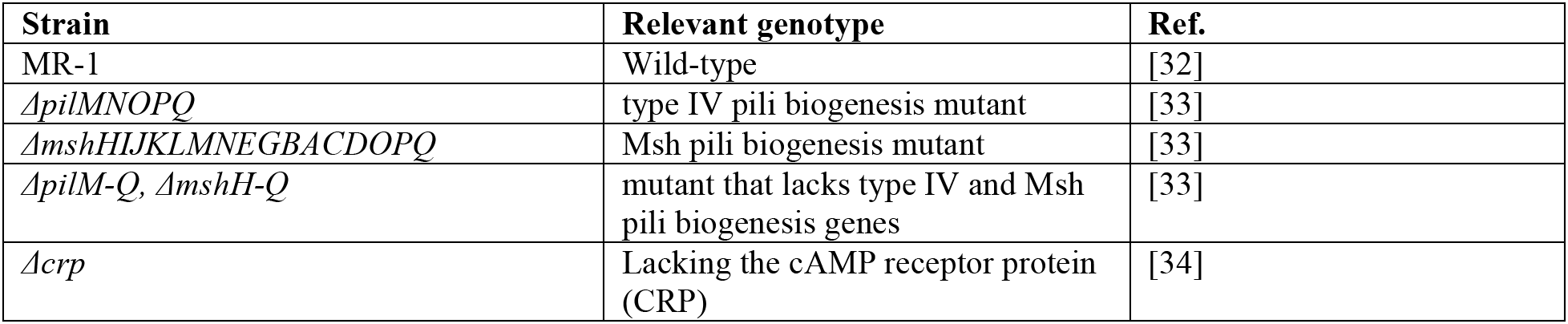
*S. oneidensis* strains used in this study

### Sample preparation for electron cryo-tomography

Cells (*L. pneumophila*, *P. aeruginosa* and *S. oneidensis*) from batch cultures and chemostats were mixed with BSA (Bovine Serum Albumin)-treated 10-nm colloidal gold solution (Sigma-Aldrich, St. Louis, MO, USA) and 4 μL of this mixture was applied to a glow-discharged, carbon-coated, R2/2, 200 mesh copper Quantifoil grid (Quantifoil Micro Tools) in a Vitrobot Mark IV chamber (FEI). Excess liquid was blotted off and the grid was plunge frozen in a liquid ethane/propane mixture for ECT imaging.

### Electron cryo-tomography

Imaging of ECT samples (*S. oneidensis* and *P. aeruginosa*) was performed on an FEI Polara 300-keV field emission gun electron microscope (FEI company, Hillsboro, OR, USA) equipped with a Gatan image filter and K2 Summit counting electron-detector camera (Gatan, Pleasanton, CA, USA). Data were collected using the UCSF Tomography software[35], with each tilt series ranging from −60° to 60° in 1° increments, an underfocus of ^~^5–10 μm, and a cumulative electron dose of ^~^130–160 e^-^/A^2^ for each individual tilt series. For *L. pneumophila* samples, imaging was done using an FEI Titan Krios 300 kV field emission gun transmission electron microscope equipped with a Gatan imaging filter and a K2 Summit direct electron detector in counting mode (Gatan). *L. pneumophila* data was also collected using UCSF Tomography software and a total dose of ~100 e^-^/A^2^ per tilt series with ~6 um underfocus.

### Sub-tomogram averaging

The IMOD software package was used to calculate three-dimensional reconstructions of tilt series[36]. Alternatively, the images were aligned and contrast transfer function corrected using the IMOD software package before producing SIRT reconstructions using the TOMO3D program[37]. Sub-tomogram averages with 2-fold symmetrization along the particle Y-axis were produced using the PEET program[38].

### Bioinformatics analysis

Candidate H- and T-ring component genes were identified by sequence alignment of the following *Vibrio cholerae* proteins against the fully sequenced genomes of each bacterial species using BLASTP. The *Vibrio cholerae* proteins used were: MotX (Q9KNX9), MotY (Q9KT95), FlgO (Q9KQ00), FlgP (Q9KQ01) and FlgT (Q9KPZ9). Candidate stator homologs in *Colwellia psychrerythraea* 34H were identified by sequence alignment of PomAB proteins of *V. cholerae* (Q9KTL0 and Q9KTK9 respectivley) and MotAB proteins of *E. coli* (P09348 and P0AF06 respectively) against its genome. The *C. psychrerythraea* 34H genome contains a single flagellar motor system[23]. Candidate MotX and MotY homologs identified were adjacent to the flagellar cluster in the genome, and for each stator system candidate homologs were characteristically located in tandem in the genome. The codes in parentheses represent Uniprot IDs. An *E*-value cutoff of < 1 × 10^-10^ was used. The raw BLAST results for all species are shown in Table S6.

## References

1. Sowa Y, Berry RM (2008) Bacterial flagellar motor. Q Rev Biophys 41.

2. Berg HC (2003) The rotary motor of bacterial flagella. Annu Rev Biochem 72: 19–54.

3. Altegoer F, Bange G (2015) Undiscovered regions on the molecular landscape of flagellar assembly. Curr Opin Microbiol 28: 98–105.

4. Kühn MJ, Schmidt FK, Eckhardt B, Thormann KM (2017) Bacteria exploit a polymorphic instability of the flagellar filament to escape from traps. Proc Natl Acad Sci 114: 6340–6345.

5. Belas R (2014) Biofilms, flagella, and mechanosensing of surfaces by bacteria. Trends Microbiol 22: 517–527.

6. Feldman M, Bryan R, Rajan S, Scheffler L, Brunnert S, Tang H, Prince A (1998) Role of flagella in pathogenesis of Pseudomonas aeruginosa pulmonary infection. Infect Immun 66: 43–51.

7. Appelt S, Heuner K (2017) The flagellar regulon of Legionella—A Review. Front Cell Infect Microbiol 7:.

8. Koebnik R (1995) Proposal for a peptidoglycan-associating alpha-helical motif in the C-terminal regions of some bacterial cell-surface proteins. Mol Microbiol 16: 1269–1270.

9. Morimoto Y, Minamino T (2014) Structure and function of the bi-directional bacterial flagellar motor. Biomolecules 4: 217–234.

10. Macnab RM (1999) The bacterial flagellum: reversible rotary propellor and type III export apparatus. J Bacteriol 181: 7149–7153.

11. Evans LDB, Hughes C, Fraser GM (2014) Building a flagellum outside the bacterial cell. Trends Microbiol 22: 566–572.

12. Kaplan M, Subramanian P, Ghosal G, Oikonomou CM, Pirbadian S, Starwalt-Lee R, Gralnick JA, El-Naggar MY, Jensen GJ (2018) Stable sub-complexes observed *in situ* suggest a modular assembly pathway of the bacterial flagellar motor. bioRxiv.

13. Oikonomou CM, Jensen GJ (2017) A new view into prokaryotic cell biology from electron cryotomography. Nat Rev Microbiol 15: 128.

14. Gan L, Jensen GJ (2012) Electron tomography of cells. Q Rev Biophys 45: 27–56.

15. Chen S, Beeby M, Murphy GE, Leadbetter JR, Hendrixson DR, Briegel A, Li Z, Shi J, Tocheva EI, Müller A, et al. (2011) Structural diversity of bacterial flagellar motors. EMBO J 30: 2972–2981.

16. Beeby M, Ribardo DA, Brennan CA, Ruby EG, Jensen GJ, Hendrixson DR (2016) Diverse high-torque bacterial flagellar motors assemble wider stator rings using a conserved protein scaffold. Proc Natl Acad Sci 113: E1917–E1926.

17. Zhu S, Nishikino T, Hu B, Kojima S, Homma M, Liu J (2017) Molecular architecture of the sheathed polar flagellum in *Vibrio alginolyticus*. Proc Natl Acad Sci 201712489.

18. Zhao X, Norris SJ, Liu J (2014) Molecular architecture of the bacterial flagellar motor in cells. Biochemistry (Mosc) 53: 4323–4333.

19. Chaban B, Coleman I, Beeby M (2018) Evolution of higher torque in Campylobacter-type bacterial flagellar motors. Sci Rep 8:.

20. Minamino T, Imada K (2015) The bacterial flagellar motor and its structural diversity. Trends Microbiol 23: 267–274.

21. Terashima H, Kawamoto A, Morimoto YV, Imada K, Minamino T (2017) Structural differences in the bacterial flagellar motor among bacterial species. Biophys Physicobiology 14: 191–198.

22. Magariyama Y, Sugiyama S, Muramoto K, Maekawa Y, Kawagishi I, Imae Y, Kudo S (1994) Very fast flagellar rotation. Nature 371: 752–752.

23. Thormann KM, Paulick A (2010) Tuning the flagellar motor. Microbiology 156: 1275–1283.

24. Doyle TB, Hawkins AC, McCarter LL (2004) The complex flagellar torque generator of Pseudomonas aeruginosa. J Bacteriol 186: 6341–6350.

25. Toutain CM, Caizza NC, Zegans ME, O’Toole GA (2007) Roles for flagellar stators in biofilm formation by Pseudomonas aeruginosa. Res Microbiol 158: 471–477.

26. Paulick A, Delalez NJ, Brenzinger S, Steel BC, Berry RM, Armitage JP, Thormann KM (2015) Dual stator dynamics in the *S hewanella oneidensis* MR-1 flagellar motor: *Shewanella oneidensis* MR-1 flagellar motor. Mol Microbiol 96: 993–1001.

27. Terashima H, Li N, Sakuma M, Koike M, Kojima S, Homma M, Imada K (2013) Insight into the assembly mechanism in the supramolecular rings of the sodium-driven Vibrio flagellar motor from the structure of FlgT. Proc Natl Acad Sci 110: 6133–6138.

28. Koerdt A, Paulick A, Mock M, Jost K, Thormann KM (2009) MotX and MotY are required for flagellar rotation in Shewanella oneidensis MR-1. J Bacteriol 191: 5085–5093.

29. Wu L, Wang J, Tang P, Chen H, Gao H (2011) Genetic and molecular characterization of flagellar assembly in Shewanella oneidensis. PLoS ONE 6: e21479.

30. Subramanian P, Pirbadian S, El-Naggar MY, Jensen GJ (2018) Ultrastructure of *Shewanella oneidensis* MR-1 nanowires revealed by electron cryotomography. Proc Natl Acad Sci 115: E3246–E3255.

31. Pirbadian S, Barchinger SE, Leung KM, Byun HS, Jangir Y, Bouhenni RA, Reed SB, Romine MF, Saffarini DA, Shi L, et al. (2014) Shewanella oneidensis MR-1 nanowires are outer membrane and periplasmic extensions of the extracellular electron transport components. Proc Natl Acad Sci 111: 12883–12888.

32. Myers CR, Nealson KH (1988) Bacterial manganese reduction and growth with manganese oxide as the sole electron acceptor. Science 240: 1319–1321.

33. Bouhenni RA, Vora GJ, Biffinger JC, Shirodkar S, Brockman K, Ray R, Wu P, Johnson BJ, Biddle EM, Marshall MJ, et al. (2010) The role of Shewanella oneidensis MR-1 outer surface structures in extracellular electron transfer. Electroanalysis 22: 856–864.

34. Charania MA, Brockman KL, Zhang Y, Banerjee A, Pinchuk GE, Fredrickson JK, Beliaev AS, Saffarini DA (2009) Involvement of a membrane-bound class III adenylate cyclase in regulation of anaerobic respiration in Shewanella oneidensis MR-1. J Bacteriol 191: 4298–4306.

35. Zheng SQ, Keszthelyi B, Branlund E, Lyle JM, Braunfeld MB, Sedat JW, Agard DA (2007) UCSF tomography: an integrated software suite for real-time electron microscopic tomographic data collection, alignment, and reconstruction. J Struct Biol 157: 138–147.

36. Kremer JR, Mastronarde DN, McIntosh JR (1996) Computer visualization of three-dimensional image data using IMOD. J Struct Biol 116: 71–76.

37. Agulleiro JI, Fernandez JJ (2011) Fast tomographic reconstruction on multicore computers. Bioinformatics 27: 582–583.

38. Nicastro D (2006) The Molecular Architecture of Axonemes Revealed by Cryoelectron Tomography. Science 313: 944–948.

